# Integrin α5β1 mediates the inhibitory effects of vasoinhibin on angiogenesis and vascular permeability

**DOI:** 10.1101/2025.07.13.664588

**Authors:** Magdalena Zamora, Carmen Clapp, Gonzalo Martínez de la Escalera, Juan Pablo Robles

## Abstract

Vasoinhibin exerts potent inhibitory effects on angiogenesis and vascular permeability through a minimal three-amino acid sequence, the HGR motif. However, the nature of the vasoinhibin receptor has remained controversial. Here, we identify integrin α5β1 as the endothelial cell-surface binding molecule mediating the actions of the HGR motif. Vasoinhibin binds to α5β1 integrin through this motif, and silencing the integrin α5 subunit abolishes the vasoinhibin-mediated inhibition of endothelial cell proliferation, invasion, and permeability. Notably, the HGR motif activates integrin α5β1, as reflected by an increase in endothelial cell adhesion to fibronectin, the canonical ligand of integrin α5β1. These findings identify integrin α5β1 as the molecular target of vasoinhibin mediating its antiangiogenic and anti-vasopermeability actions. Furthermore, a novel integrin activation mechanism leading to suppressed angiogenesis is unveiled, thereby challenging the conventional integrin inhibition approach as a therapeutic intervention.

## Introduction

Angiogenesis, the formation of new blood vessels, is essential during development and becomes largely quiescent in adulthood (1). In pathological conditions such as cancer and diabetic retinopathy, angiogenesis becomes exacerbated and directly contributes to disease progression (2). Angiogenesis results from the complex interplay of diverse molecular and cellular pathways (3), involving stimulators such as vascular endothelial growth factor (VEGF) (4), and inhibitors, including peptide hormones (5–7). Notably, many of these inhibitors are proteolytic fragments of proteins not originally associated with angiogenesis (8), such as vasoinhibin.

Vasoinhibin is a naturally occurring fragment of the hormone prolactin that potently inhibits angiogenesis and vascular permeability by suppressing angiogenic signaling pathways, including those triggered by VEGF (9, 10). Beyond its antiangiogenic effects, vasoinhibin promotes fibrinolysis, endothelial cell apoptosis, and inflammation (10). The molecular mechanisms that enable vasoinhibin to exert its diverse biological effects remain poorly understood. Previous studies have identified two distinct membrane molecular partners for vasoinhibin: the multimeric complex formed by plasminogen activator inhibitor-1 (PAI-1), urokinase plasminogen activator (uPA), and urokinase receptor (uPAR), which was proposed to mediate its antiangiogenic and fibrinolytic properties (11); and integrin α5β1, associated with its pro-apoptotic effects (12). These findings imply that different structural determinants within vasoinhibin mediate its binding to these specific molecular partners.

We recently identified two distinct functional motifs in the vasoinhibin molecule: the HGR and the HNLSSEM motifs. However, the biological effects of these structural determinants do not align with the actions of the previously reported molecular binding partners of vasoinhibin. Although the HNLSSEM motif mediates vasoinhibin binding to PAI-1, this interaction promotes fibrinolysis, inflammation, and apoptosis, rather than inhibition of angiogenesis and vascular permeability (13). Moreover, while the HGR motif potently inhibits angiogenesis and vascular permeability (14), it does not bind PAI-1 (13). These discrepancies raised the need to explore the possibility that integrin α5β1 may be the molecular target for the HGR motif to inhibit angiogenesis and vascular permeability.

Integrin α5β1 belongs to a large family of heterodimeric cell-surface receptors composed of α and β subunits, which mediate adhesion to the extracellular matrix and activate signaling pathways regulating cell growth, migration, and survival (15). Integrin α5β1 and its canonical ligand, fibronectin, are particularly critical for angiogenesis, as their coordinated expression in endothelial cells forms a provisional extracellular matrix necessary for vessel formation (16–18). Specifically, α5β1 integrin recognizes the RGD and PHSRN motifs in fibronectin, facilitating endothelial cell anchoring and triggering signaling cascades leading to the angiogenesis process (17, 19, 20). Despite the essential role of this integrin in angiogenesis, its therapeutic blockade has shown limited clinical success, primarily due to unexpected agonist-like activities exhibited by some antagonists and the simplistic assumption that mere integrin α5β1 inhibition should be therapeutically sufficient (21, 22). These challenges have underscored the complexity of integrin-targeted therapies and emphasized the need for additional therapeutic approaches beyond simple integrin disruption.

This work demonstrates that the HGR motif binds to integrin α5β1 and that this interaction mediates the potent inhibition of angiogenesis and vascular permeability exerted by vasoinhibin. Furthermore, binding to the HGR motif did not inhibit integrin α5β1-mediated adhesion, but instead, it enhanced endothelial cell adhesion to fibronectin. These results encourage the clinical translation of HGR-based peptides and introduce novel integrin-activating approaches, rather than conventional inhibitory strategies, for suppressing angiogenesis.

## Results

### Vasoinhibin binds to integrin α5β1 through its HGR motif

To evaluate whether the HGR motif of vasoinhibin binds to integrin α5β1, we treated human umbilical vein endothelial cells (HUVEC) with a biotinylated peptide comprising residues 45 to 51 of vasoinhibin (B-Vi45-51), which includes the HGR motif, and performed a pull-down assay. The lysates were incubated with streptavidin magnetic beads, followed by Western blot analysis of the previously reported vasoinhibin binding partners. Immunoreactive integrin subunit α5 was precipitated with B-Vi45-51 more than with either the non-biotinylated Vi45-51 or the streptavidin beads alone, and corresponded to a ∼145 kDa protein, consistent with its reported molecular mass (Figure 1A). Treatment with VEGF and bFGF did not alter the binding profile. We also evaluated whether B-Vi45-51 could pull down PAI-1 or uPAR, the other described vasoinhibin binding partners (11), but no immunoreactivity was detected for either protein.

**Figure 1.**
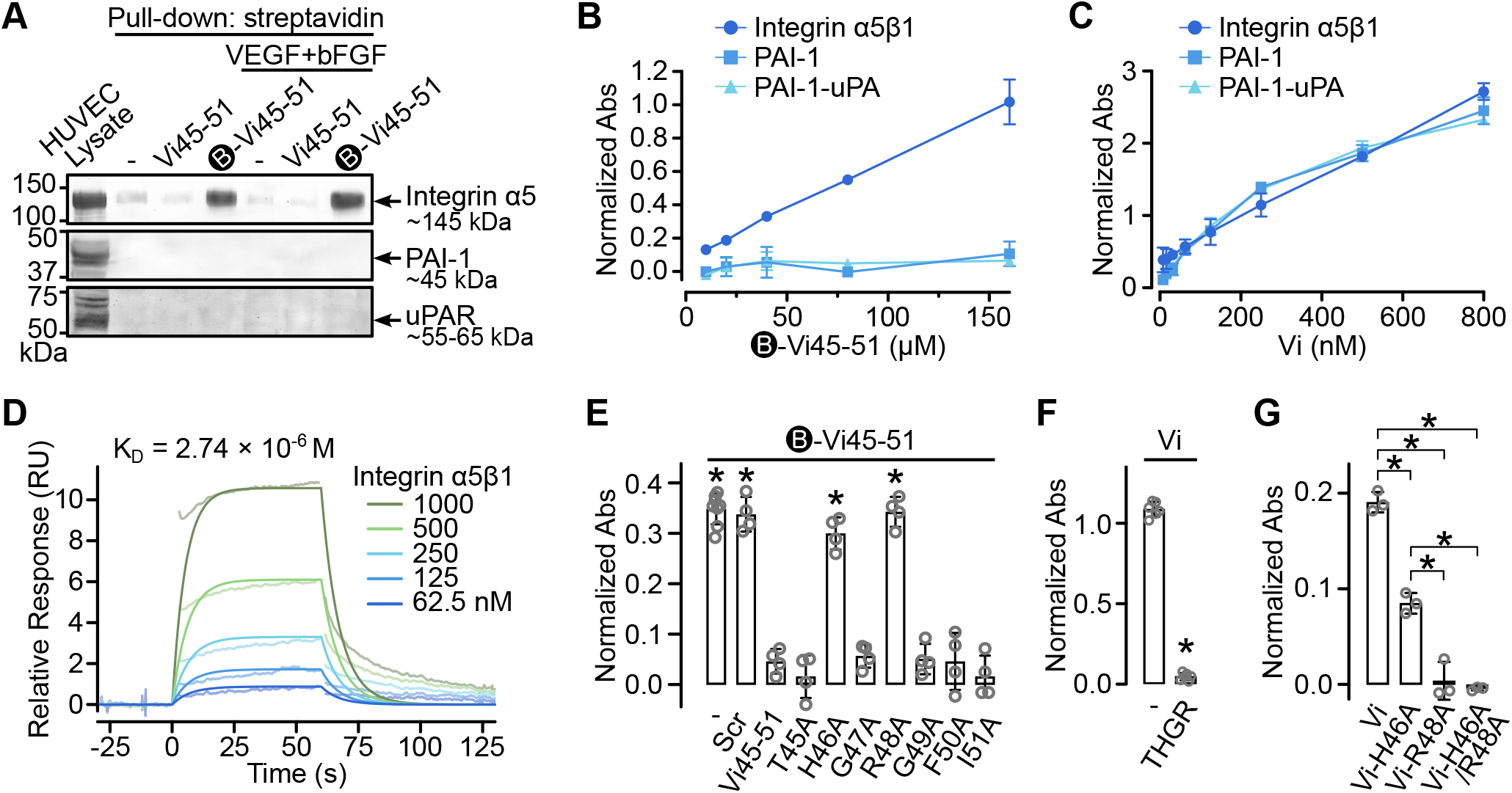
Binding analysis of the HGR motif to integrin α5β1. (**A**) Representative pull-down-Western blot assay of HUVEC lysates treated without (-) or with Vi45-51 or biotinylated Vi45-51 (B-Vi45-51) peptides in the absence or presence of VEGF+bFGF and blotted with antibodies against integrin α5, PAI-1, and uPAR. Expected molecular masses are indicated. ELISA evaluating dose-dependent binding of B-Vi45-51 (**B**) or full-length vasoinhibin (Vi) (**C**) to immobilized integrin α5β1, PAI-1, or the PAI-1-uPA complex. (**D**) Surface plasmon resonance (SPR) analysis of integrin α5β1 binding to immobilized B-Vi45-51 at various concentrations. Binding affinity and kinetics were calculated using a steady-state 1:1 binding model. (**E**) Competitive ELISA evaluating interference of B-Vi45-51 binding to immobilized integrin α5β1 with scrambled peptide (Scr), Vi45-51, or alanine-scanning mutants. \**P*<0.001 vs. Vi45-51 (one-way ANOVA, Dunnett). (**F**) Competitive ELISA assay evaluating interference of Vi binding to immobilized integrin α5β1 using the HGR-containing tetrapeptide THGR. \**P*<0.001 vs. Vi (unpaired t-test). (**G**) ELISA assessing binding to integrin α5β1 of Vi and its alanine mutants Vi-H46A, Vi-R48A, or Vi-H46A/R48A. \**P*<0.001 (one-way ANOVA, Tukey). Individual points are independent replicates. Data represent mean ± SD of at least three independent experiments.

Consistent with the pull-down results, an ELISA demonstrated a dose-dependent binding of B-Vi45-51 to immobilized integrin α5β1, but not to immobilized PAI-1 or the PAI-1-uPA complex (Fig. 1B). In contrast, full-length vasoinhibin bound to integrin α5β1, PAI-1, and the PAI-1-uPA complex (Fig. 1C), with calculated equilibrium dissociation constant (K_D_) values of 383.2 nM, 527.9 nM, and 405.8 nM, respectively, like previously reported affinities (11). Surface plasmon resonance (SPR) analysis further confirmed the direct interaction of integrin α5β1 with the immobilized B-Vi45-51, and a K_D_ of 2.74 μM (Fig. 1D) consistent with reported affinities for integrin interactions with small peptide motifs (23). These findings support the hypothesis that vasoinhibin binds integrin α5β1 via its HGR motif.

To confirm that the HGR motif is responsible for integrin α5β1 binding, we performed competitive ELISAs evaluating the interaction between immobilized integrin α5β1 and B-Vi45-51. Non-biotinylated Vi45-51 inhibited this interaction, whereas a heptapeptide with a scrambled sequence had no effect. Alanine-scanning mutations further demonstrated that substituting either H46 or R48 with alanine (H46A or R48A) in Vi45-51 abolished competitive inhibition (Fig. 1E), highlighting these residues as critical for integrin binding. Moreover, the tetrapeptide THGR entirely blocked the interaction between vasoinhibin and integrin α5β1, confirming integrin recognition via the HGR motif (Fig. 1F). Consistently, alanine mutants of recombinant vasoinhibin (Vi-H46A, Vi-R48A, and Vi-H46A/R48A), demonstrated reduced integrin binding; specifically, Vi-H46A partially reduced binding (∼60%), while Vi-R48A and the double mutant Vi-H46A/R48A completely abolished it. These findings strongly support the essential role of the HGR motif in integrin α5β1 recognition.

### Integrin α5β1 is required for the antiangiogenic activity of the HGR motif

To evaluate whether integrin α5β1 is necessary for the antiangiogenic activity of the HGR motif, we knocked down the integrin α5 subunit gene (*ITGA5*) in HUVECs using lentiviral-delivered shRNA (shITGA5). This resulted in a 92.5% reduction in *ITGA5* mRNA (Fig. 2A) and a corresponding 91.5% decrease in integrin α5 protein (Fig. 2B). Functionally, *ITGA5* silencing reduced HUVEC adhesion to fibronectin, the canonical integrin α5β1 ligand, by approximately 50% (Fig. 2C). The adhesion of *ITGA5*-deficient cells to non-fibronectin-coated surfaces was unaffected, confirming effective and specific inhibition of integrin α5β1. Since integrin α5 only pairs with β1 (24), other integrins were expected to remain unaffected. Compared to uncoated conditions, HUVECs showed significantly greater adhesion (>40%) to fibronectin-coated plates. The incomplete reduction in fibronectin binding by shITGA5 likely reflects compensation by other integrins (15).

**Figure 2.**
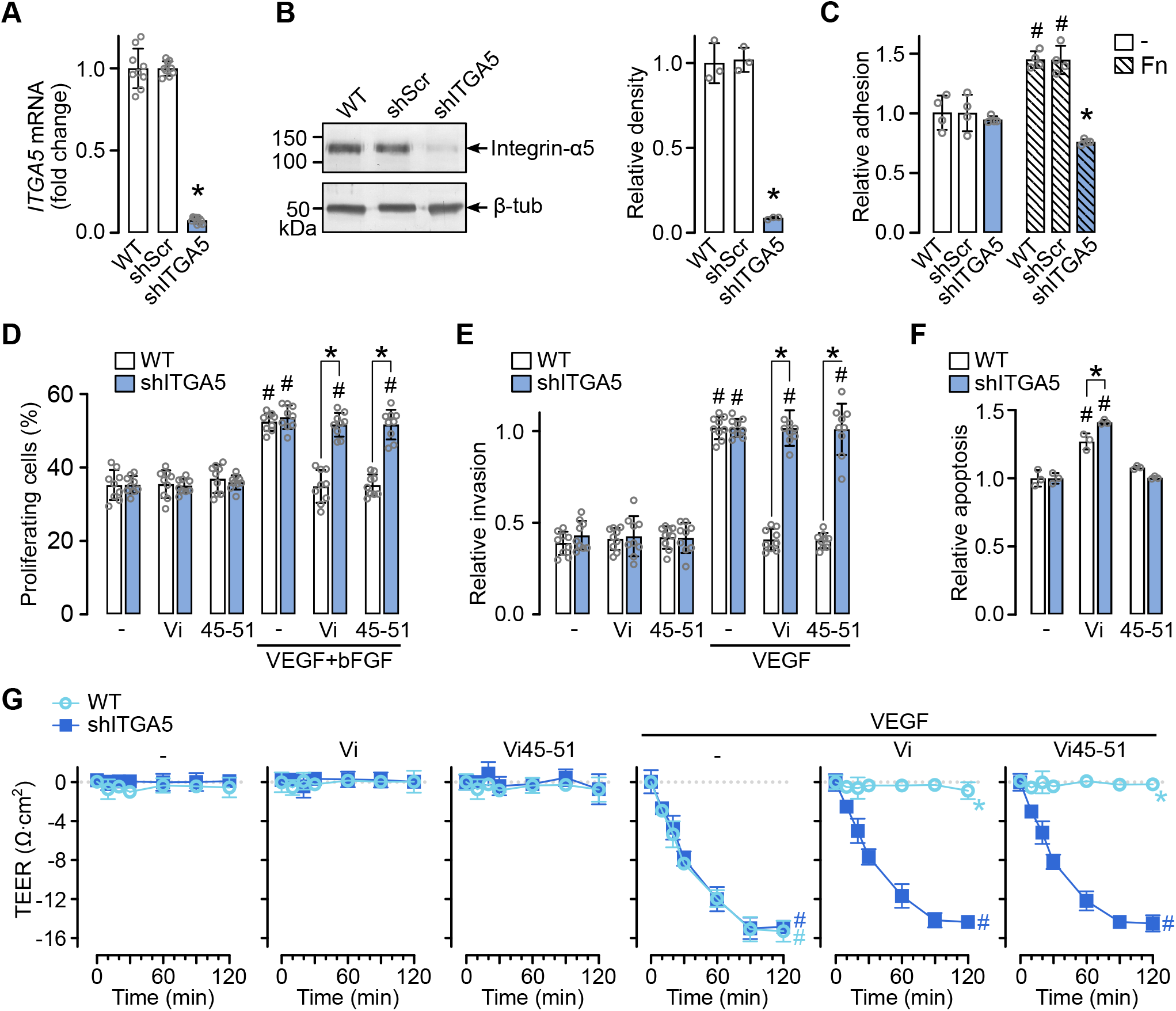
Effect of integrin α5 knockdown on the antiangiogenic activity of the HGR motif. (**A**) mRNA expression levels of *ITGA5* in wild-type (WT) HUVEC or transduced with lentiviral scrambled (shScr) control or targeting integrin α5 (shITGA5) shRNA. n=9, \**P*<0.001 (one-way ANOVA, Dunnett). (**B**) Representative Western blot and densitometric quantification of integrin α5 (MW 145 kDa) in lysates of WT, shScr, and shITGA5 HUVEC lysates. β-tubulin (50 kDa). n=3, \**P*<0.001 (one-way ANOVA, Dunnett). (**C**) Adhesion of WT, shScr, or shITGA5 HUVEC to uncoated or fibronectin-coated plates (Fn). n=4, #*P*<0.001 vs. -, \**P*<0.001 vs. WT (two-way ANOVA, Sidak and Dunnett). Proliferation (**D**) and matrigel invasion (**E**) of WT and shITGA5 HUVEC treated with full-length vasoinhibin (Vi) or Vi45-51, stimulated by VEGF+bFGF or VEGF. n=9, #*P*<0.001 vs. unstimulated cells, \**P*<0.001 vs. WT (two-way ANOVA, Sidak and Dunnett). (**F**) Apoptosis of WT and shITGA5 HUVEC treated with Vi or Vi45-51, n=3, #*P*<0.001 vs. -, \**P*=0.003 vs. WT (two-way ANOVA, Sidak and Dunnett). (**G**) Transendothelial electrical resistance (TEER) measurements of WT or shITGA5 HUVEC monolayers in transwell inserts treated with Vi or Vi45-51, in the absence or presence of VEGF. n=6, #*P*<0.001 vs. absence of VEGF, \**P*<0.001 vs. WT (repeated measures ANOVA). Individual points are independent replicates. Data represent mean ± SD of at least three independent experiments.

Knockdown of *ITGA5* completely abolished the inhibitory effects of both vasoinhibin and Vi45-51 on endothelial cell proliferation and invasion (Fig. 2D, E), demonstrating that integrin α5β1 mediates the antiangiogenic properties of the HGR motif. Likewise, the inhibitory effect of vasoinhibin and Vi45-51 on the VEGF-induced vascular permeability was completely lost in *ITGA5*-deficient cells (Fig. 2G). *ITGA5* knockdown does not affect basal endothelial proliferation, migration, permeability, or the response to stimulation with VEGF and bFGF.

Since it has been previously reported that vasoinhibin induces endothelial apoptosis through the integrin α5β1 (12), we evaluated HUVEC apoptosis in *ITGA5*-deficient cells. *ITGA5* knockdown did not abolish vasoinhibin-induced apoptosis; in fact, it was even slightly increased (Fig. 2F), indicating that integrin α5β1 is not responsible for the proapoptotic effect of vasoinhibin. This is consistent with the previous report that the HNLSSEM motif, through uPAR, mediates apoptosis induced by vasoinhibin (13). These findings show that integrin α5β1 mediates the antiangiogenic and anti-permeability effects of the HGR motif of vasoinhibin.

### The HGR motif does not inhibit integrin α5β1-mediated adhesion

Since integrin α5β1 inhibition is known to suppress angiogenesis by disrupting endothelial cell adhesion to the extracellular matrix (17), we assessed whether the HGR motif similarly disrupted endothelial cell adhesion. As expected, the RGD tripeptide, which interferes with the adhesion of many integrins (25), inhibited HUVEC adhesion to fibronectin-coated plates in a dose-response manner (IC_50_ = 61.29 nM). Unexpectedly, Vi45-51 increased the adhesion in a dose-dependent manner (EC_50_ = 4.55 nM) (Figure 3A). In contrast, a scrambled version of the Vi45-51 had no effect on adhesion. Furthermore, an HGR tripeptide increased adhesion like Vi45-51, but the Vi45-51 carrying an R48A mutation did not (Figure 3B), confirming the role of the HGR motif in enhancing endothelial adhesion mediated by the fibronectin receptor, integrin α5β1. To further confirm this distinct mechanism, we compared Vi45-51 with known integrin inhibitors in this adhesion assay, including the RGD-mimetic cilengitide (26), the PHSRN-based peptide ATN-161 (27), and the monoclonal antibody volociximab (28). As expected, all the inhibitors suppressed adhesion, while Vi45-51 further enhanced the fibronectin-induced cell adhesion of endothelial cells (Figure 3C).

**Figure 3.**
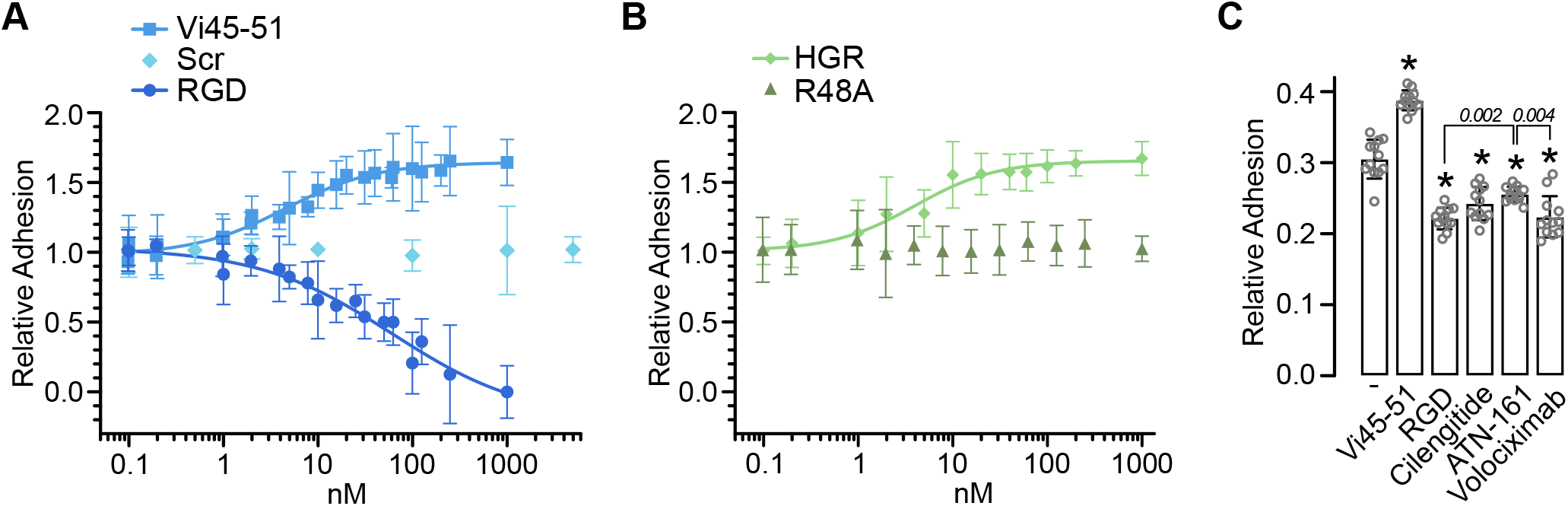
Effect of the HGR motif on integrin α5β1-mediated endothelial cell adhesion. Dose-response adhesion of HUVECs onto fibronectin-coated plates in the presence of Vi45-51, scrambled (Scr), or RGD tripeptide (**A**), or HGR tripeptide and Vi45-51 mutated by alanine in R48 (R48A) (**B**). Curves represent nonlinear regression fits (*r*^*2*^ >0.8). (**C**) Adhesion of HUVECs treated with 1 μM of integrin inhibitors (cilengitide, ATN-161, and volociximab) on fibronectin-coated plates. \**P* < 0.001 vs. -, (one-way ANOVA, Tukey). Individual points are independent replicates. Data represent mean ± SD of at least three independent experiments.

## Discussion

Integrins have emerged as promising therapeutic targets due to their significant roles in pathological processes, particularly in exacerbated angiogenesis. However, integrin-targeted therapies have thus far shown limited clinical success (22). In this study, we identified integrin α5β1 as the molecular target responsible for mediating the potent inhibitory effects of the vasoinhibin HGR motif on angiogenesis and vascular permeability (Figure 4). Interestingly, contrary to the prevailing therapeutic paradigm seeking integrin inhibition, the HGR motif activated integrin α5β1 to downregulate angiogenesis, thereby unveiling an alternative modulatory mechanism by which integrin-targeted therapies could operate.

**Figure 4.**
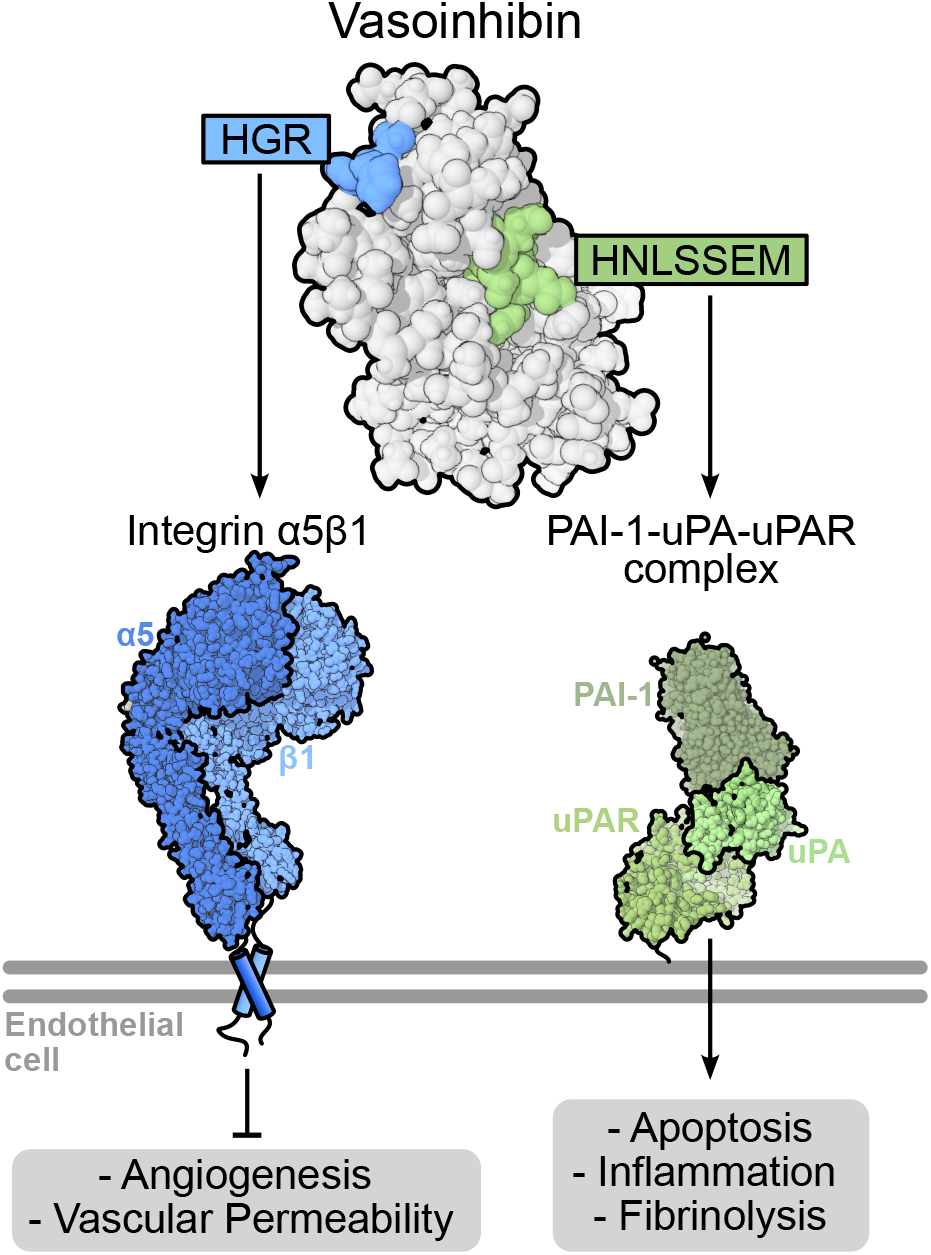
Vasoinhibin contains two distinct motifs that independently regulate vascular function through specific molecular targets. The HGR motif of vasoinhibin binds to integrin α5β1 to inhibit angiogenesis and vascular permeability. This interaction also enhances endothelial adhesion to fibronectin. In contrast, the HNLSSEM motif mediates the binding of vasoinhibin to the PAI-1-uPA-uPAR complex, inducing endothelial apoptosis, inflammation, and fibrinolysis.

The HGR motif is the sole antiangiogenic determinant of vasoinhibin, as its mutation leads to the loss of vasoinhibin’s antiangiogenic function (14). Given that vasoinhibin binds to integrin α5β1 via its HGR motif and that knockdown of ITGA5 abolishes the effects of vasoinhibin on endothelial cell proliferation, invasion, and vascular permeability, integrin α5β1 emerges as the binding partner mediating the antiangiogenic activity of vasoinhibin. Integrin α5β1 is critical for angiogenesis (16) and is linked to endothelial cell signaling pathways known to be downregulated by vasoinhibin, such as the PI3K/Akt and MAPK pathways (10). However, the precise molecular signaling mechanisms downstream of the vasoinhibin–integrin α5β1 interaction remain unknown, and our study cannot exclude the contribution of additional molecules, distinct from but influenced by integrin α5β1. Along this line, vasoinhibin binds to other endothelial cell surface proteins, specifically the PAI-1– uPA–uPAR multimeric complex (11). The antiangiogenic and antitumor effects of vasoinhibin are lost in PAI-1 knockout mice, implying that binding to PAI-1 is essential for vasoinhibin’s actions (11). This concept is further supported by reports showing a functional link between integrin α5β1 and uPAR, as their interactions critically modulate endothelial cell migration, extracellular matrix remodeling, and intracellular signaling pathways during angiogenesis (29). Nevertheless, the HGR motif itself does not bind PAI-1 (present data, 13), and peptides containing this motif retain potent antiangiogenic properties without PAI-1 binding (13, 14). Moreover, PAI-1 is known to be proangiogenic (30), and its genetic deletion alone impairs angiogenesis (11). Also, the PAI-1–uPA– uPAR complex lacks intracellular domains to directly transduce antiangiogenic signals (29). And while these observations clearly exclude PAI-1 as the mediator of vasoinhibin antiangiogenic properties, binding to PAI-1 via the HNLSSEM motif accounts for inflammatory, apoptotic, and fibrinolytic actions of vasoinhibin (Figure 4) and might indirectly influence the regulation of angiogenesis within physiological contexts such as tissue repair (31), a possibility awaiting investigation.

Given the combinatorial complexity inherent to the different structural determinants of vasoinhibin, it is not surprising that different conclusions have been drawn regarding its key biological mediators. Another example is integrin α5β1. Harigaya’s group first described the binding of vasoinhibin to integrin α5β1 and showed that antibodies against integrin α5β1 partially prevented vasoinhibin-induced endothelial cell apoptosis (12), concluding that integrin α5β1 mediates the apoptotic effects of vasoinhibin. In contrast, we found no evidence to support this claim. Silencing integrin α5β1 did not prevent, but slightly enhanced, endothelial cell apoptosis induced by vasoinhibin, indicating that integrin α5β1 is not involved in its apoptotic mechanism. This discrepancy may be explained by the fact that certain integrin α5β1 antibodies can actively promote cell survival rather than acting as passive blockers (32). Integrin inhibitors such as cilengitide (26), RGD and ATN-161 peptides (27), or monoclonal antibody volociximab (28), which disrupt endothelial adhesion, induce apoptosis through anoikis (33). However, the HGR motif enhances, rather than disrupts, endothelial cell adhesion to fibronectin, and peptides containing this motif are not proapoptotic (13). Furthermore, our findings align with the previously described HNLSSEM motif of vasoinhibin, which induces endothelial apoptosis through a uPAR-dependent mechanism (13). Altogether, these observations support the conclusion that the integrin α5β1 is not the mediator of the apoptotic properties of vasoinhibin.

The unexpected observation that the HGR motif enhances, rather than inhibits, endothelial cell adhesion to fibronectin represents a regulatory mechanism distinct from conventional integrin-blocking strategies. Integrin-mediated adhesion typically promotes the proliferation and survival of endothelial cells (34, 35), which is why inhibitory therapeutic compounds (e.g., cilengitide, ATN-161, volociximab, endostatin, and tumstatin) have been developed as antiangiogenic therapies (26–28). However, under certain contexts, integrin-mediated adhesion may inhibit cell proliferation and migration. For instance, the absence of β3, β5, or α3β1 integrins in endothelial cells enhances angiogenesis (36, 37). Moreover, increased cell adhesion may hinder proliferation, as endothelial cells forming capillary tubes in fibronectin-rich matrices exhibit growth arrest (38). Further evidence of integrin-mediated growth suppression emerges from oncology research. Overexpression of integrin α5β1 can reduce the proliferation of cancer cells (39–43), and cancer cells that acquire anchorage-independent proliferation typically downregulate integrin α5β1 (44). This functional duality underscores the complexity inherent to integrin regulation of cellular processes.

The mechanism by which increased integrin α5β1-mediated endothelial adhesion suppresses angiogenesis warrants further investigation. Given the notable difference between the potencies of the HGR motif to induce adhesion (EC_50_ ∼4.5 nM) and to inhibit proliferation (IC_50_ ∼150 pM) of endothelial cells, with a relatively low K_D_ (2.74 μM) suggest that these effects might not only result from mechanical cues, such as cellular shape, tension, motility, or matrix stiffness (45). Instead, this disparity suggests the involvement of finely tuned signaling amplification pathways downstream of integrin α5β1 activation. Potential mechanisms might include modulation of Rho/Rac signaling components (46), blockage of kinases that regulate cell-cycle progression (42), downregulation of critical growth factor receptors (43), or lateral inhibitory interactions with other cell surface receptors (47).

In conclusion, this study identifies integrin α5β1 as the molecular target for the antiangiogenic effects of vasoinhibin HGR motif and highlights a promising novel mechanism of integrin modulation. These insights advance our understanding of integrin biology and may open therapeutic avenues beyond conventional integrin-inhibitory strategies.

## Experimental procedures

### Materials

Peptides including Vi45-51, biotinylated Vi45-51 (B-Vi45-51), scrambled control, single-residue mutants (T45A, H46A, G47A, R48A, G49A, F50A, I51A), HGR, and THGR were synthesized by GenScript (Piscataway, NJ). RGD, Cilengitide, and ATN-161 were from Sigma-Aldrich (St. Louis, MO); Volociximab from Novus Biologicals (Centennial, CO). Recombinant human integrin α5β1was from Bio-Techne (Minneapolis, MN), and fibronectin from Gibco (Waltham, MA). Recombinant vasoinhibin (123 residues) (48) and ViH46A, ViR48A, and ViH46A/R48A (14) were produced as reported. Human umbilical vein endothelial cells (HUVEC) (49) were cultured in F12K with 20% fetal bovine serum, 100 μg mL^-1^ heparin (Sigma-Aldrich), and 25 μg mL^-1^ ECGS (Corning, NY). VEGF (GenScript); bFGF donated by Scios, Inc. (Mountain View, CA). Antibodies for Western blot included integrin α5 (Sigma-Aldrich), uPAR (Bio-Techne), β-tubulin, PAI-1 (Abcam, Cambridge, UK), and respective secondary antibodies.

### Pull-Down Assay and Western Blot

HUVECs were incubated with 500 nM biotinylated Vi45-51 (B-Vi45-51), 20 ng mL^-1^ bFGF, and 25 ng mL^-1^ VEGF for 1 h. Cells were lysed in RIPA, sonicated, and incubated at 37ºC for 30 min. Pull-down used Dynabeads Streptavidin T1 (Thermo-Fisher, Waltham, MA). Eluted proteins were resolved by SDS-PAGE, transferred to nitrocellulose membrane, and probed for integrin α5, PAI-1, and uPAR using AP secondary antibodies.

### Enzyme-Linked Immunosorbent Assay (ELISA)

Plates coated with 2 µg mL^-1^ integrin α5β1, PAI-1, or PAI-1-uPA (pre-incubated at 37ºC, 10 min) were incubated with different concentrations of B-Vi45-51 or vasoinhibin, or 100 nM ViH46A, ViR48A, or ViH46A/R48A. Competition assays used excess scrambled peptide, alanine mutants, or THGR. Detection used HRP-conjugated antibodies and TMB substrate.

### Surface Plasmon Resonance (SPR)

B-Vi45-51 (100 nM) was immobilized on a streptavidin-coated SA chip (Biacore 1K, Cytiva) at 10 μL min^-1^. Integrin α5β1 (62.5-1000 nM) was injected (30 µL min^-1^, 60 s association/dissociation) in HBS-P buffer (150 mM NaCl, pH 7.4, and 1 mM MnCl_2_). Kinetics were evaluated using a 1:1 binding model. Chip regeneration was done in running buffer.

### shRNA-mediated Integrin α5 Knockdown

HUVECs were transduced with pLKO.1-based MISSION shRNA vectors targeting ITGA5 sequence 5’-CCTCAGGAACGAGTCAGAATT-3’ (TRCN0000029652) (NM_002205,Sigma-Aldrich) or scrambled control #1864 (Addgene). Knockdown was confirmed by qPCR and Western blot.

### Endothelial Cell Proliferation

HUVEC wild-type (WT) or ITGA5-silenced (shITGA5) cells were seeded in 96-well plates, serum-starved (0.5% FBS) for 8 h, and then treated for 24 h with 100 nM Vi45-51 or vasoinhibin plus bFGF and VEGF in 20% FBS. As reported, EdU incorporation was detected with Azide Fluor 545 (Sigma-Aldrich) (14).

### Invasion

HUVECs (WT and shITGA5) were seeded on Matrigel-coated Transwells (0.38 mg mL^-1^) in starvation medium with 100 nM of Vi45-51 or vasoinhibin. 3T3-L1 conditioned medium with 50 ng mL^-1^ VEGF served as chemoattractant. After 16 h, invading cells were fixed, stained, and counted.

### Apoptosis

HUVECs (WT/shITGA5) were seeded at 13,000 cells cm^-2^ in 12-well plates. After serum starvation (0.5% FBS, 4h), cells were treated with 100 nM vasoinhibin or Vi45-51 for 24 h. Floating and adherent cells were collected, adjusted to 100,000 cells mL^-1^, and analyzed using the Cell Death Detection ELISA kit (Roche, Basel, CH).

### Endothelial Permeability

WT/shITGA5 HUVECs were grown to confluency on 0.4 μm pore Transwells, treated with 100 nM Vi45-51 or vasoinhibin for 1 h, then stimulated with 50 ng mL^-1^ VEGF. Trans-endothelial electrical resistance (TEER) was recorded for 120 min using an EVOM2 meter (World Precision Instruments, Sarasota, FL).

### Endothelial Adhesion

96-well plates, coated or not with fibronectin (1 μg cm^-2^), were blocked with 20% FBS-F12K and incubated with Vi45-51, HGR, R48A, scrambled, or RGD, Cilengitide, ATN-161, or Volociximab. HUVECs (43,000 cells cm^-2^) were added and incubated at 37°C for 1h. Cells were washed twice, incubated with 0.5 mg mL^-1^ MTT for 3 h, added DMSO, and absorbance measured at 570 nm.

### Statistical analysis

One-way ANOVA followed by the Dunnett or Tukey post hoc test, unpaired t-tests, two-way ANOVA followed by Sidak or Dunnett post hoc tests, and repeated-measures ANOVA were performed using GraphPad Prism version 10.5.0 for MacOS (GraphPad Software, San Diego, California, USA). The overall significance threshold was set at *P* < 0.05.

## Data Availability

All data generated or analyzed during this study are included in this article.

## Acknowledgments

We thank Xarubet Ruíz Herrera and Fernando López Barrera for their technical assistance.

## Author contributions

MZ, CC, GME, and JPR conceptualization: MZ and JPR methodology, MZ investigation, CC, GME, and JPR supervision, GME funding acquisition, CC, and JPR writing-original draft, and MZ, CC, GME, and JPR writing-review & editing.

